# Aβ-targeting synNotch Receptor for Alzheimer’s Disease: Expanding Applications to Extracellular Protein Aggregates

**DOI:** 10.1101/2024.10.15.618096

**Authors:** Nicholas J. Bergo, Suckwon Lee, Cynthia J. Siebrand, Julie K. Andersen, Chaska C. Walton

## Abstract

The synthetic Notch receptor (synNotch) system is a versatile platform that induces gene transcription in response to extracellular signals. However, its application has been largely confined to membrane-bound targets due to specific activation requirements. Whether synNotch can also target extracellular protein aggregates, such as amyloid beta (Aβ) in Alzheimer’s disease (AD), is unclear. To address this, we engineered an Aβ-targeting synNotch receptor controlling the production of chimeric human-mouse versions of Lecanemab (Leqembi®) or Aducanumab (Aduhelm®), both FDA-approved antibodies for AD. We demonstrate that NIH 3T3 cells expressing this synNotch system detect and respond to extracellular Aβ aggregates by synthesizing and secreting Aducanumab or Lecanemab. These findings broaden the potential applications of synNotch, extending its targets beyond membrane-bound proteins to extracellular protein aggregates, providing obvious benefits to research in this scientific arena.

## Introduction

Recently, the FDA has approved three immunotherapies for Alzheimer’s disease (AD)— Aducanumab (Aduhelm®), Donanemab (Kisunla™), and Lecanemab (Leqembi®)—all targeting amyloid-beta (Aβ), offering a promising means to slow cognitive and functional decline associated with AD^1–4^. Although many additional therapeutics are currently being tested in clinical trials^5^, largely the ability to halt or significantly slow the disease has not been achieved.

A mostly unexplored alternative for new interventions or improving existing ones in the AD field is synthetic biology. Synthetic modular receptors present an exciting opportunity for the development of novel targeted cell-based therapies^6^. A well-known example is the chimeric antigen receptor (CAR), used in cytotoxic CD8 T-cells (CAR-T) for cancer treatment^7^. CARs link extracellular cues to native immunomodulatory functions via endogenous signaling cascades. There are additional synthetic receptors that can link extracellular cues to transcriptional programs using orthogonal signaling^6^. In this case, the receptors do not activate native cell functions. Instead, they allow for custom sense-and-respond programs that can trigger the expression of artificial genes. A well-established example is the synthetic Notch receptor (synNotch)^8,9^.

One major application of synNotch receptors is to regulate the expression of CARs to enhance specificity and efficacy in CAR-T cell therapies^10–14^. Alternatively, synNotch can be engineered to create “sense-and-respond” biological drug factories. In cancer treatment research, synNotch receptors have been used to trigger the synthesis and secretion of therapeutic agents, such as antibodies or cytokines, in response to specific antigens on target cells^9,15^. Although still unexplored, the use of synNotch for precision delivery of therapeutic peptides directly to the brain could be a promising approach to minimize the side effects of traditional bulk drug delivery methods, such as intravenous injection.

A key factor in designing synthetic receptors is their antigen-binding domain (ABD), which determines the antigen that activates the receptor^6,7^. In cancer applications, the ABD of synNotch receptors mainly targets antigens present on the membrane of cells^6^. In contrast, AD is characterized by extracellular protein aggregates, particularly Aβ plaques^16,17^. A novel application of synNotch receptors would be to target these extracellular Aβ aggregates, making them ideal candidates for delivering Aβ-clearing antibodies like Lecanemab. However, synNotch receptors are not activated by soluble monomers unless tethered to the cell membrane^6,8,18^. Whether this limitation extends to non-membrane-bound aggregates, such as Aβ plaques, remains unknown. Gaining insight into this would be crucial for expanding synNotch applications to AD and other diseases involving similar extracellular aggregates.

To answer this, we present tests conducted in NIH 3T3 cells using a receptor system developed in our laboratory that integrates synNotch and CAR technology for cell-based drug delivery^19^. This system includes an Aβ-targeting synNotch receptor based on Aducanumab (Adu-synNotch), designed to regulate the expression of murine versions of Lecanemab and Aducanumab. Our results demonstrate that Adu-synNotch is activated by extracellular Aβ aggregates, leading to the synthesis and secretion of the therapeutic antibodies.

## Results & Discussion

To assess the feasibility of targeting extracellular Aβ aggregates with synNotch receptors, we developed an *in vitro* model of AD in which we treated Adu-synNotch-expressing NIH 3T3 cells with Aβ (Figure 1a). Aducanumab was chosen to template the antigen-binding domain of synNotch because of its broader selectivity for Aβ aggregate species when compared to other Aβ-targeting antibodies^1,4,20^. The antigen binding domain consisted of a single chain variable fragment (scFv) with the amino acid sequences of the variable light chain (VLC) and variable heavy chain (VHC) of Aducanumab fused by a flexible G4S(3) linker (Figure 1b, Supplementary Figure 1a). The rest of the sequence of the synNotch receptor has been described previously^8,9^.

**Figure 1.**
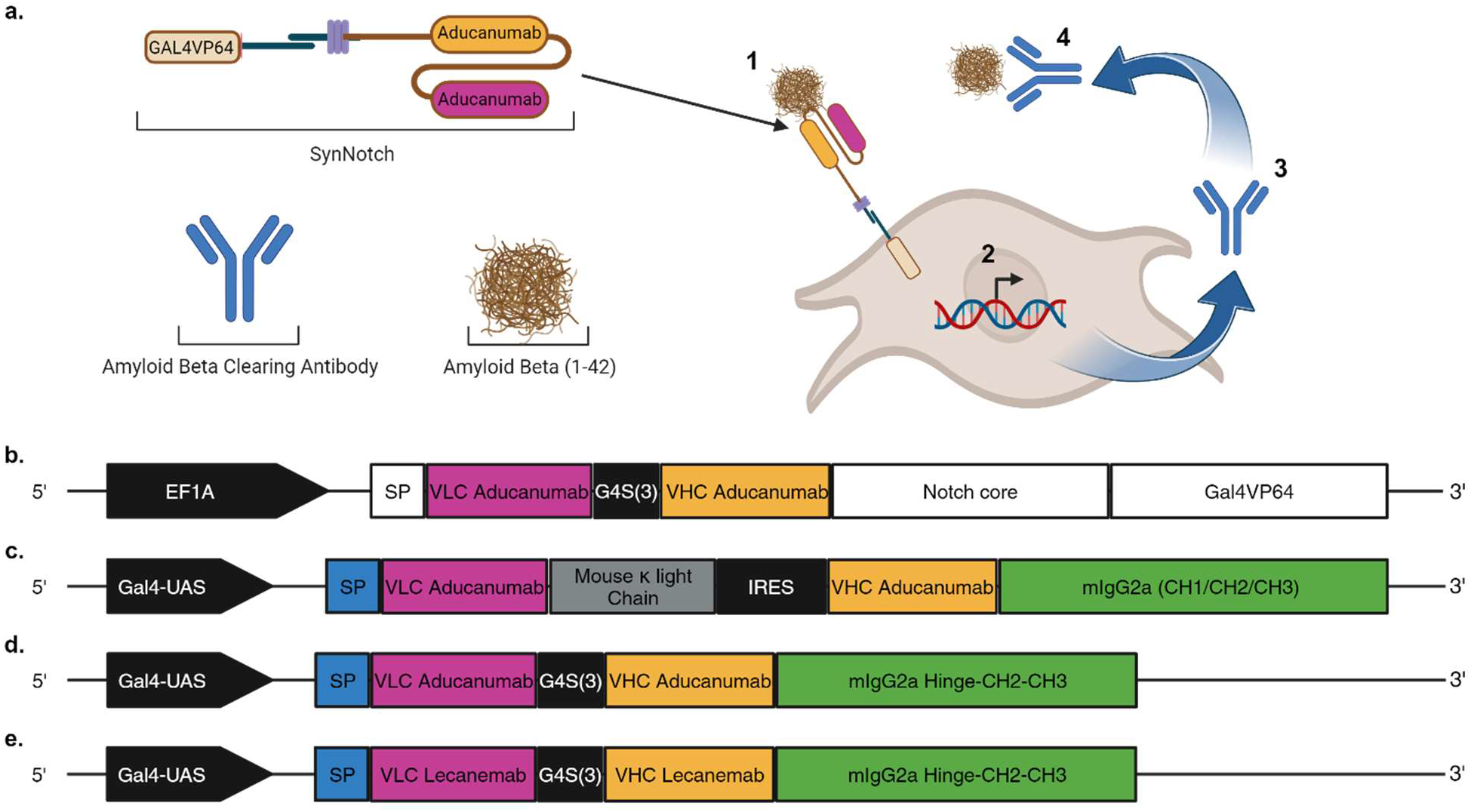
AD model to assess the ability of Aβ-targeting synNotch to be activated and induce production of a therapeutic payload. **(a)** Adu-synNotch regulating a therapeutic antibody expressed in NIH 3T3 cells. **1**. Treatment of cells with Aβ preparations. **2**. Activation of Adu-synNotch results in the transcription of an artificial gene coding a therapeutic antibody. **3**. Secretion of the antibody. **4**. Binding of the antibody to Aβ. (**b**) Expression cassette for EF1A-regulated Adu-synNotch. The VLC and VHC of Aducanumab are fused by a flexible G4S(3) linker to form an scFv, with an N-terminal human CD8α signal peptide (SP; white) for membrane localization. (**c**) chAducanumab antibody containing the VLC of Aducanumab fused to a mouse kappa light chain (Uniprot; P01837). Separated by an IRES, the VHC of Aducanumab fused to the constant heavy chain and Fc of mouse IgG2a (Uniprot; P01863). (**d**) scFv-Fc Aducanumab: scFv Aducanumab fused to the hinge of mouse IgG2a. (**e**) scFv-Fc chLecanemab: scFv containing VLC and VHC of Lecanemab with a G4S(3) linker, fused to the hinge of mouse IgG2a. (**c-e**) Secrecon secretory signal peptide (SP; blue).

Several constructs were tested as the therapeutic payloads to be regulated by Adu-synNotch receptors. First, we generated a human-mouse chimeric Aducanumab (chAducanumab) antibody that was previously used in mice^4^. Aducanumab itself is a human IgG1 monoclonal antibody isolated from cognitively normal healthy aged individuals with selectivity for aggregated Aβ oligomers (AβO)^4^. chAducanumab consists of a fragment antigen-binding (Fab) region formed by the variable regions of human Aducanumab fused to mouse constant chains and a fragment crystallizable (Fc) region based on a mouse heavy chain (Figure 1c, Supplementary Figure 1b)^4^. In addition to chAducanumab Fab-Fc, we generated a shorter chAducanumab scFv-Fc version (Figure 1d; Supplementary Figure 1c) as well as a chimeric human-mouse Lecanemab (chLecanemab) scFv-Fc (chLecanemab; Figure 1e; Supplementary Figure 1d). Lecanemab is a humanized IgG1 monoclonal antibody developed to target Aβ protofibrils^21^.

To assess Adu-synNotch response to Aβ, we used mixed Aβ_(1-42)_ aggregate preparations (Aβagg). These consisted of mixed AβO 0.1 µM plus Aβ fibrils (AβF) 0.5 µM enriched preparations, obtained from incubating Aβ_(1-42)_ for 3 hours or 7 days, respectively (Supplementary Figure 2). NIH 3T3 cells, either untransduced or transduced with Adu-synNotch, were treated with a mixture of AβO 0.1 µM plus AβF 0.5 µM. We performed time-lapse experiments to follow chAducanumab production in live cells. For these experiments, chAducanumab or chLecanemab were identified by a monoclonal mouse IgG2a secondary antibody, which was added to the live cell cultures. Importantly, the latter meant that non-specific antibody binding could not be blocked. Time-lapse videos initiated 2 hours after addition of the Aβagg cocktail showed no production of chAducanumab in untransduced cells (Video 1) whilst Adu-synNotch-expressing cells showed a clear increase (Video 2). We performed additional experiments in which we added the conjugated pan-Aβ antibody 6E10 and the mouse IgG2a secondary antibody to live cells. These time-lapse videos initiated after 6 hours of treatment with Aβagg plus antibodies. For untransduced cells, time-lapse images show Aβagg with clear 6E10 immunostaining and only sporadic non-specific signal on the chAducanumab channel (6E10, Video 3; chAducanumab, Video 4). In contrast, Adu-synNotch expressing cells showed both 6E10 and clear chAducanumab signal (6E10, Video 5; chAducanumab, Video 6).

We next performed immunostaining for fibrillar Aβ with OC antibody for colocalization with chAducanumab^23^. Adu-synNotch-expressing cells were treated with either vehicle (VEH) or Aβagg. Two days later, a secondary anti-mouse IgG2a antibody was added to live cells to track the progression of chAducanumab production, and cells were fixed and blocked for OC immunocytochemistry (ICC) after two more days. As above, blocking of non-specific binding could not be performed when adding secondary anti-mouse IgG2a to live cells. Immunostaining showed no chAducanumab signal colocalized with OC in VEH-treated cells (Figure 2a). Presence of non-specific staining for chAducanumab was observed but not for OC. Cells treated with Aβagg, however, showed strong OC signal colocalized with chAducanumab (Figure 2a).

**Figure 2.**
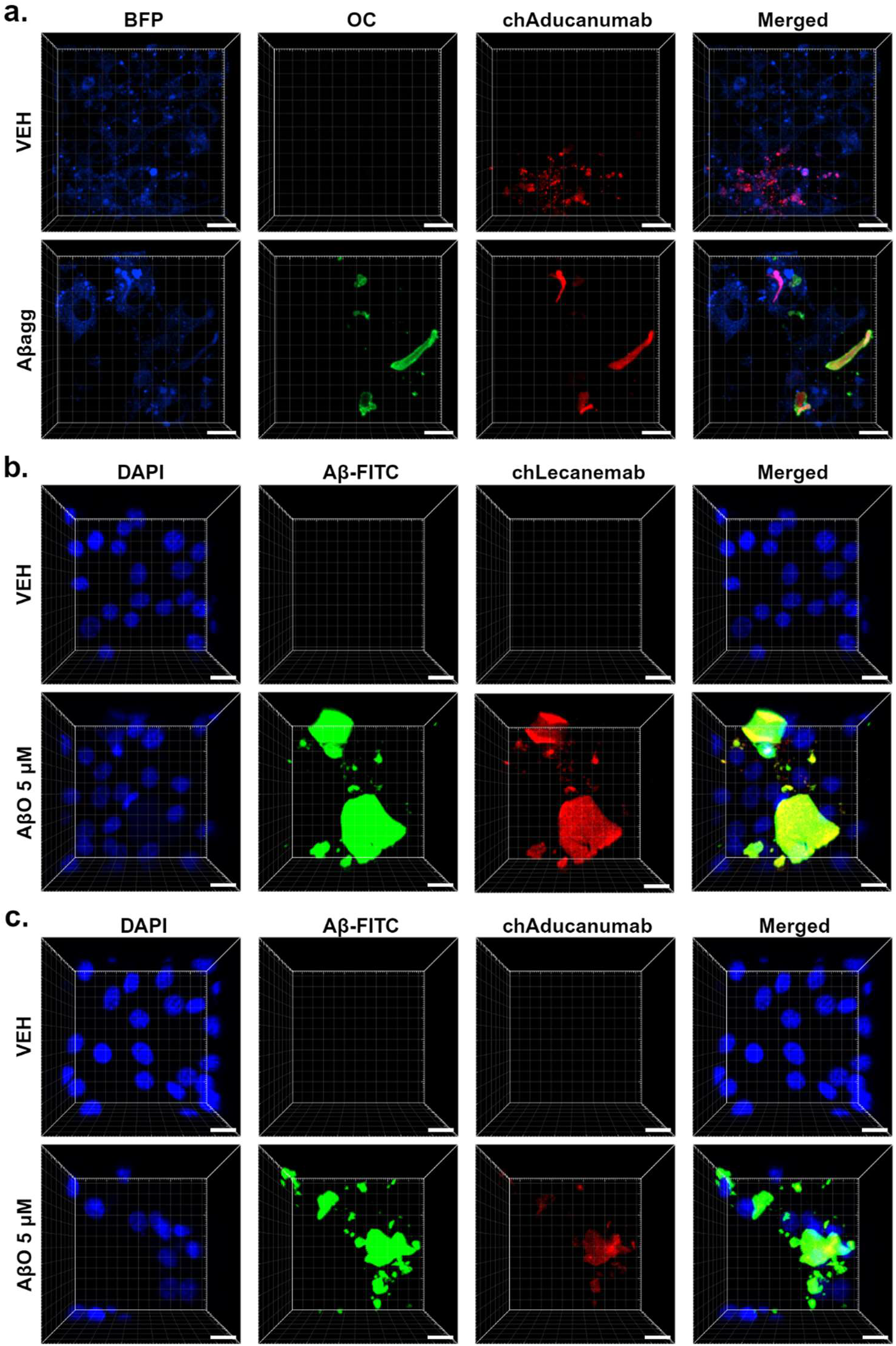
Adu-synNotch cells produce chAducanumab and chLecanemab in response to Aβ_(1-42)_ treatments. **(a)** Representative 3D confocal images of NIH 3T3 clones expressing Adu-synNotch regulating a Fab-Fc chAducanumab. Cells were treated with AβO 0.1 µM plus AβF 0.5 µM cocktail (Aβagg) for 4 days and fixed for ICC with anti-fibrillar Aβ antibody OC. To identify live production of chAducanumab, mouse IgG2a secondary antibody was added to live cells 2 days after Aβagg. Blocking of non-specific binding was possible for OC ICC (fixed cells) but not for mouse IgG2a (live cells), explaining non-specific immunostaining of chAducanumab in VEH. Blue fluorescent protein (BFP) identifies Adu-synNotch-expressing cells. Representative images of one experiment. **(b, c)** Representative 3D confocal images of NIH 3T3 clones expressing Adu-synNotch regulating either scFv-Fc chLecanemab (**b**), or scFv-Fc chAducanumab (**c)** 48 hours after treatment with VEH or AβO, 5 µM. Aβ aggregates were identified by a FITC tag, cell nuclei by DAPI, and chAducanumab/chLecanemab by anti-mouse IgG2a secondary antibody. Representative images of three independent experiments. Scale bar: 20 µm.

Lecanemab is currently favored over Aducanumab, due to slightly better results in clinical trials^1^. This motivated us to also make and test an Adu-synNotch system regulating chLecanemab. To assess Adu-synNotch response to Aβ, we used Fluorescein (FITC) tagged Aβ_(1-42)_ (Aβ-FITC). Aβ-FITC was incubated at 37° C for 3 hours to obtain preparations enriched in AβO. NIH 3T3 cells were treated with oligomer-enriched Aβ-FITC preparations (AβO-FITC) at 5 µM or VEH for 48 hours, at which time they were fixed for ICC. Cells were identified with DAPI, and chAducanumab or chLecanemab were immunostained with a monoclonal mouse IgG2a secondary antibody. For VEH-treated cells, confocal microscope images revealed no Aβ-FITC, chLecanemab, or chAducanumab signal (Figure 2b, c). In contrast, both chLecanemab (Figure 2b) and chAducanumab (Figure 2c) colocalized with Aβ-FITC aggregates under 5 µM AβO-FITC-enriched treatment conditions.

Given that chLecanemab was demonstrated to be produced by Adu-synNotch-expressing cells and is arguably the clinically favored intervention, we performed further quantification using this construct. For these experiments, NIH 3T3 cells were treated with oligomer-enriched AβO-FITC preparations as described above at 0.05 µM, 0.5 µM, and 5 µM or with VEH. After 48 hours, cells were fixed for ICC. Confocal microscope images showed no Aβ-FITC nor chLecanemab signal under VEH treatment conditions (Figure 3a). Control untransduced NIH 3T3 cells treated with 5 µM AβO did not show chLecanemab signal in any of the replicates (Supplementary Figure 3a). As expected, Aβ-FITC signal was shown to be significantly different between treatment conditions, *F*(2, 6) = 56.363, *p* < .001, *ω*^2^ = .949 (Supplementary Figure 3b). Aβ-FITC increase between 0.05 and 0.5 µM AβO treatments was not significant (p = .922), but there was a significant increase between 0.5 and 5 µM AβO treatments (p < .001) (Supplementary Figure 3b). As a positive control, Aβ-FITC aggregates were positive for the anti-APP/Aβ antibody 6E10 (Supplementary Figure 3c). A clear signal for chLecanemab was readily evident in 0.5 µM and 5 µM AβO treatment conditions (Figure 3a). Quantification of mean chLecanemab signal within confocal images revealed a statistically significant difference between treatment conditions *F*(2, 6) = 17.524, *p* = .003, *ω*^2^ = .854 (Figure 3b). Unless otherwise specified, descriptive statistics are means and standard deviations. The mean signal intensity for chLecanemab for AβO at 0.05, 0.5 and 5 µM were 2123.41 (± 505.74), 12503.99 (± 3450.19), and 6852.33 (± 1308.54), respectively.

**Figure 3.**
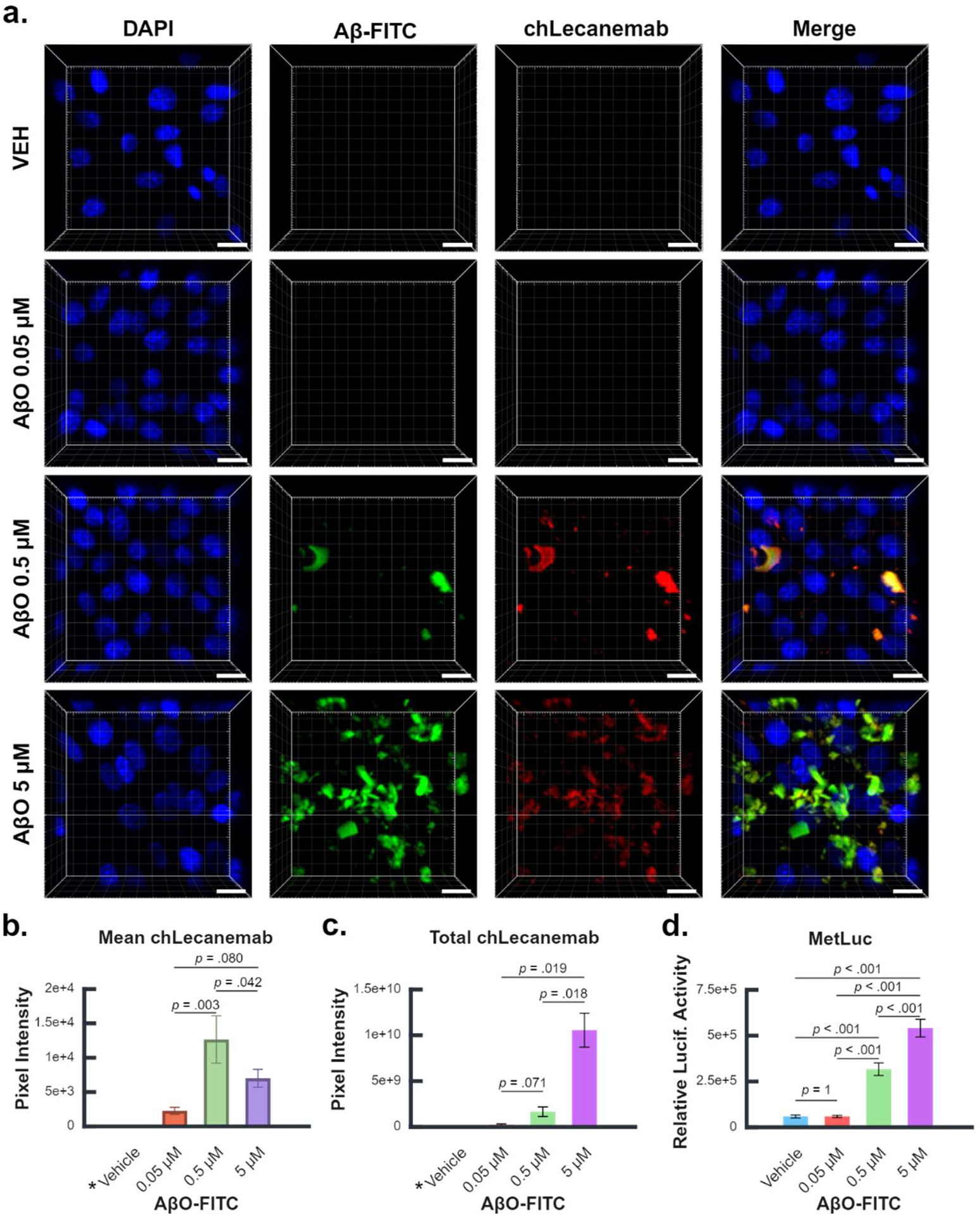
chLecanemab produced by Adu-synNotch-expressing cells binds Aβ-FITC aggregates. **(a)** Representative 3D confocal images of NIH 3T3 clones expressing Adu-synNotch 48 hours after VEH, 0.05, 0.5, or 5 µM AβO-FITC treatments with mouse IgG2a immunostaining of chLecanemab. Aβ aggregates are identified by a FITC tag and cell nuclei by DAPI. FITC and chLecanemab are readily visible at 0.5, or 5 µM AβO-FITC treatments. Representative images of three independent experiments. (**b**) Quantification of mean chLecanemab signal colocalized with FITC. Multiple comparisons with Tukey’s honestly significant difference (HSD). (**c**) Quantification of total (sum) chLecanemab signal colocalized with FITC. Multiple comparisons with Games Howell Post-hoc tests. (**b, c**) *Vehicle FITC-chLecanemab colocalization was not quantified due to lack of detectable FITC signal. **(d)**. Quantification of relative Luciferase activity in the media of treated cells. Multiple comparisons with Tukey’s HSD. (**b-d**) Results are from three independent experiments. Error Bars 1x SE. Scale bar: 20 µm.

Tukey multiple comparisons revealed an increase in chLecanemab signal between 0.05 µM and 0.5 µM AβO preparations (10380.5851, 95% CI, 4993.4821 to 15767.6880, *p* = .003) and a significant decrease from 0.5 µM and 5 µM AβO treatments (-5651.6521, 95% CI, -11038.7551 to -264.5491, *p* = .042). An increase in mean signal intensity from 0.05 µM to 5 µM (4728.9329, 95% CI, -658.1700 to 10116.0359) displayed a non-significant trend (*p* = .08). This data indicates that mean amount of chLecanemab per Aβ aggregate is larger for AβO treatments at 0.5 µM than at 5 µM, potentially reflecting that chLecanemab being produced by cells at 0.5 µM is enough to saturate Aβ aggregates whilst the amount being produced at 5 µM is not.

The sum signal for chLecanemab reflects the total amount of antibody bound to Aβ aggregates, which can shed light on the total production of chLecanemab by NIH 3T3 cells. The quantification of sum chLecanemab signal in confocal images revealed a statistically significant difference between treatment conditions, Welch’s *F*(2, 2.757) = 45.684, *p* = .008 (Figure 3c). The mean sum signal of chLecanemab for AβO-FITC preparations at 0.05, 0.5 and 5 µM treatments were 1.26 × 10^8^ (± 9.47 × 10^7^), 1.57 × 10^9^ (± 5.25 × 10^8^), and 1.05 ×10^10^ (± 1.85 × 10^9^), respectively. Games-Howell multiple comparisons revealed that the increase between 0.05 µM and 5 µM (1.03 ×10^10^, 95% CI, 4.05 × 10^9^ to 1.66 × 10^10^), *p* = .019) and the increase between 0.5 µM and 5 µM (8.90 ×10^9^, 95% CI, 3.24 × 10^9^ to 1.46 × 10^10^, *p* = .018) AβO-FITC treatments were statistically significant, while the increase between 0.05 µM and 0.5 µM (1.42 × 10^9^, 95% CI, -2.78 × 10^8^ to 3.12 × 10^9^) preparations only displayed a trend that was not statistically significant (*p* = .071) (Figure 3c). This data demonstrates that more total chLecanemab is bound to Aβ aggregates with increasing AβO concentrations.

In the experiments described above, an undetermined amount of chLecanemab could have remained in the media if the amount of antibody produced by NIH 3T3 cells treated with 0.05 µM and 0.5 µM AβO-FITC preparations was high enough to saturate Aβ aggregates. To accurately assess synNotch activation, we included a secreted Metridia Luciferase (MetLuc) expressed in-frame with chLecanemab and separated by a *Thosea asigna* virus 2A (T2a) sequence with an N-terminal added Furin cleavage site and a GSG linker (Supplementary Figure 1d)^24–26^. MetLuc also enabled the quantification of Adu-synNotch ligand-independent activation (LIA) under VEH treatment conditions. MetLuc luminescence was found to be significantly different between treatment conditions, *F*(3, 8) = 179.44, *p* < .001, *ω*^2^ = 0.985 (Figure 3d). Means for VEH, AβO 0.05, AβO 0.5 µM, and AβO 5 µM treatments were 53555 (± 8048), 53500 (± 6117), 311681 (± 34583), and 535823 (± 48136), respectively. Tukey post hoc analysis revealed that luminescence of AβO 0.05 µM was 55 (95% CI, -78663 to 78553) units less than for VEH, but this was a non-statistically significant difference (*p* = 1.000). However, the increase between VEH and AβO 0.5 µM treatment (258126, 95% CI, 179518 to 336733), VEH and AβO 5 µM (482268, 95% CI, 403660 to 560876), AβO 0.05 µM and AβO 0.5 µM (258180, 95% CI, 179573 to 336788), AβO 0.05 µM and AβO 5 µM (482323, 95% CI, 403715 to 560931), and AβO 0.5 µM and AβO 5 µM (224142, 95% CI, 145535 to 302750) were all highly statistically significant (*p* < .001). This MetLuc analysis revealed that Adu-synNotch receptors displayed a dose-response activation to extracellular AβO beginning at a concentration of AβO 0.5 µM. Adu-synNotch production of MetLuc at AβO 5 µM was higher than that observed at AβO 0.5 µM, evidencing a dose-dependent synNotch response. Cells treated with AβO 0.05 µM, however, were not different from VEH, likely reflecting synNotch LIA, as already reported by others^8,27,28^.

In summary, we provide the first *in vitro* proof-of-concept for a synNotch-based intervention for AD. Our results show synNotch can target extracellular Aβ aggregates and deliver Aβ-clearing antibodies. Expression of synNotch receptors in appropriate carrier cell systems would provide a means of direct delivery of therapeutic payloads to brain regions burdened by toxic proteinaceous aggregates, including Aβ. These results are envisioned to be extendable to other diseases characterized by proteinaceous aggregates.

## Acknowledgments

Funding was provided by NIH RF1 AG068296 & R01 AG081989 (JKA, CCW, SL, NB). The figures were prepared using BioRender.

## Author Contributions

N.J.B. and S.L. performed the experiments. Confocal Imaging was done by N.J.B., S.L., and C.C.W. S.L. made the cell clones with the sleeping beauty system and the western blot. C.J.S. contributed to the design of the experiments and produced the cell clones with lentivirus. Image processing and ELISA was done by N.J.B. C.C.W. designed the constructs and performed statistical analysis. J.K.A. and C.C.W. supervised and guided the project. N.J.B. and C.C.W. wrote the manuscript, with input from all other authors.

## Declaration of Interests

J.K.A., C.C.W., and S.L. are inventors in Patent No. WO2024076500A3.

## Methods

### Constructs

The Sleeping Beauty Transposon transposase system was used for experiments with Adu-SynNotch and chAducanumab Fab-Fc (Figure 1b, c)^22^. Adu-synNotch expression was regulated from an EF-1α core promoter. chAducanumab Fab-Fc was regulated by an inducible Gal4UAS mini-CMV sequence described previously^8^. An additional expression cassette included a CAR, a tagBFP reporter protein, and a Blasticidin (BSD) antibiotic resistance gene regulated by a human PGK promoter. The 2A sequences included a Furin cleavage site and a GSG linker to minimize the C-terminal amino acid additions after 2A-induced ribosomal skipping^26^. CARs and additional genes are part of a larger system being developed by our laboratory to provide clinical AD interventions^19^. These constructs were not eliminated from the plasmids described above and the transfer vectors described below because synNotch receptors operate through an orthogonal signaling pathway that does not interfere with and is not interfered by endogenous signaling^8,9^. The constructs were expressed from a pT4-HB transposon plasmid. pT4/HB was a gift from Wolfgang Uckert (Addgene, Plasmid #108352; http://n2t.net/addgene:108352; RRID:Addgene_108352)^29^. Expression cassettes were codon optimized for expression in mice, synthesized, subcloned, and sequence-verified by Genescript. SP1-SB100X transposase was used for transposition. SP1-SB100X was a gift from Joseph Dougherty & Rob Mitra (Addgene, Plasmid #154887; http://n2t.net/addgene:154887; RID:Addgene_154887)^30^.

Lentiviral delivery was used to transduce Adu-synNotch regulating chAducanumab scFv-Fc or chLecanemab scFv-Fc (Figure 1b, d, e). The synNotch receptor and payload (chAducanumab or chLecanemab plus MetLuc) were expressed from two separate third-generation self-inactivating (SIN) lentiviral vectors based on pLV[Exp]-EGFP/Puro-EF1A>mCherry transfer vector (VectorBuilder, VB010000-9298rtf). SynNotch expression was regulated from an EF1A promoter. Downstream of the synNotch receptor, there was a murine Foxp3 (mFoxp3) gene and a SNAP reporter protein, separated by T2A and P2A sequences, respectively, with Furin and GSG sequences as described above. chAducanumab and chLecanemab, as well as MetLuc, are regulated by a Gal4UAS mini-CMV sequence as described above. This vector included a second expression cassette regulating the expression of a CAR and an mNeonGreen reporter protein separated by a T2A sequence. All vectors were codon optimized for expression in mice, synthesized, subcloned, and sequence-verified by VectorBuilder.

### Cell Culture, lipofection, and transduction

NIH/3T3 mouse fibroblast adherent cells were purchased from ATCC (ATCC^®^, CRL-1658^™^). NIH/3T3 cells were cultured without antibiotics in DMEM (VWR, 45000-304) supplemented with 10% FBS (VWR, 45000-734) at 37°C, 5% CO2.

For lipofection of sleeping beauty plasmids, NIH/3T3 cells were seeded in Ibidi 8-well slides (Ibidi; 80826) using DMEM supplemented with 10% FBS and incubated overnight. Lipofectamine™ 3000 (Thermo Fisher Scientific, L3000008) was used for lipofection according to the manufacturer instructions. Lipoplexes were prepared at a ratio of 1 μl of Lipofectamine per 1 μg of DNA. Each well was transfected with 0.6 µg of transposon carrying Adu-synNotch and chAducanumab and 0.06 µg of SP1-SB100X transposase (10:1 transposon:transposase ratio). Transfected clones were selected for BSD resistance by treating cells with 10 µg/mL BSD (Invivogen, Ant-bl-05) for 7 days. BSD-resistant cells were frozen in DMEM supplemented with 10% FBS and 5% DMSO. For lentiviral transduction, NIH/3T3 cells were seeded in Ibidi 8-well slides using the DMEM supplemented with 10% FBS and incubated overnight. NIH/3T3 cells were co-transduced at a 1000 MOI (by physical titer) with 5 µg/mL polybrene transduction enhancer (VectorBuilder, PL0001). Transduction media was replaced 24 hours later with fresh DMEM supplemented with 10% FBS. 48 hours later, cells were incubated with SNAP-cell SiR 647 substrate (NEB, S9102S) and Live Dead Fixable Violet (Invitrogen, L34955), as recommended by the manufacturers. Following SNAP substrate incubation, double-positive clones were sorted by SNAP+/mNeonGreen+ markers using a BD FACS Aria II cell sorter. Cells were expanded and sorted a second time to guarantee high purity of SNAP+/mNeonGreen+ expression. Cells were frozen in DMEM supplemented with 10% FBS and 5% DMSO.

### Aβ treatments

Amyloid beta: Beta amyloid (1-42) aggregation kit was purchased from rPeptide (rPeptide, A-1170-2). Lyophilized Aβ peptides were reconstituted in 5mM Tris or 10mM NaOH at a concentration of 1mg/ml and diluted with HPLC water and Tris-buffered saline (TBS) to obtain a 100 µM stock solution. Preparations to obtain solutions enriched in oligomer and solutions enriched in fibrils were done as previously described, with some modifications^31^. To obtain solutions enriched in oligomers or fibrils, the stock solution was incubated for 3 hours at 37°C or for 7 days at 37°C, respectively. For Aβ treatments, cell clones were thawed per the vendor’s instructions (ATCC^®^, CRL-1658^™^) and passaged prior to Aβ treatments. Approximately 6×10^4^ cells were plated per well into treated µ-Slide 8 Well chambers (Ibidi, 80826) and allowed to adhere overnight. Cells were then incubated with Aβ preparations at the specified durations and concentrations. FITC-labeled Amyloid beta: Lyophilized FITC labeled Aβ42 peptides (rPeptide, A-1119-1) were reconstituted in 50 µL DMSO (0.5 mg in 50 µL), diluted 1:10 in Neurobasal Plus culture medium (Gibco, A3582901) and incubated at 37°C for 3 hours. VEH consisted of 50 µL of DMSO diluted in Neurobasal Plus medium (Gibco, A3582901). For AβO-FITC treatments, cell clones were thawed per the vendor’s instructions and passaged prior to Aβ treatments. Approximately 6×10^3^ sorted double-positive SNAP and mNeonGreen NIH/3T3 clones were seeded per well into treated µ-Slide 8 Well chambers and allowed to adhere overnight. Cells were then incubated with AβO-FITC enriched preparations at the specified concentrations for 48 hours, after which supernatants were collected for luciferase assay and the cells were fixed for ICC.

### Immunofluorescence and microscopy

Immunostaining and confocal imaging were performed without removing cells from µ-Slide 8 Well chambers. For ICC, cells were fixed for 10 minutes with 4% paraformaldehyde (Santa Cruz, 3052589-4) at room temperature (RT), followed by three 10-minute washes with ice-cold PBS and incubated with 10% NGS (Jackson ImmunoResearch, 005-000-121) in PBS for 30 min at RT to block unspecific antibody binding. Cells were then incubated for 1 hour at RT with primary antibodies in 10% NGS in PBS. After 4 washes with PBS, cells were incubated for 1 hour at RT with secondary antibodies in 10% NGS in PBS. After 4 washes with PBS, DNA labeling was performed using PBS containing 1 µg/ml 4′,6-diamidino-2-phenylindole (DAPI) (1:10000) (Invitrogen, D1306). After 3 additional washes with PBS, PBS was completely replaced by pure glycerol (Invitrogen, 15514-011). For live imaging, antibodies were directly added to the media. Confocal images, tile scan images, and live tile scan images were obtained with the Zeiss LSM 780 Confocal Microscope using an EC Plan-Neofluar 20X/0.50 M27 objective. Live cell imaging was performed in a sealed chamber at 37 °C and 5% CO2.

### Western Blot

Standard (Bio-Rad, 1610376) and 6.8 µg of amyloid beta samples with fluorescent compatible sample buffer (Invitrogen, LC2570) were loaded into the 4-12% NuPAGE Bis-Tris Gel (Invitrogen, NP0321BOX). The gel was run with NuPAGE MES SDS running buffer (Invitrogen, NP0002) at 50V for 30 mins for stacking, and 100V for separation. After the run, PVDF was activated with methanol for 1 min and rinsed with the 1x transfer buffer (25 mM Tris, 192 mM Glycine, and 20% v/v methanol). The transfer was run at 25V for 2 hours. PVDF was washed with PBS/0.1% Tween 20 (PBST) (VWR, 0777-1L) three times for 5 mins. PVDF was blocked with 10% NGS in PBST for 1hr at RT and washed with PBST. Anti-amyloid beta 6E10 in PBST was incubated for 1 hour at RT and washed 3 times with PBST. Secondary antibody anti-mouse IgG1 AF488 (1:3000) (Invitrogen, A-21121) was added and incubated for 1 hour at RT and washed 3 times with PBST. The membrane was imaged by ChemiDoc MP Imaging System (Bio-Rad).

### Antibodies

Primary antibodies: anti-amyloid fibril OC antibody (1:100, ICC) (Sigma-Aldrich, AB2286), anti-β-amyloid 1-16 6E10 (1:1000, WB) (Biolegend, 8030001). Secondary antibodies: anti-rabbit IgG AF488 (1:500, ICC) (Invitrogen, A-11008), anti-mouse IgG1 AF488 (1:3000, WB) (Invitrogen, A-21121), and anti-mouse IgG2a AF546 (1:500) (Invitrogen, A-21133) for chAducanumab and chLecanemab for ICC and live imaging. Conjugated antibodies: Alexa Fluor 594 conjugated anti-β-amyloid 1-16 6E10 (1:200) (Biolegend, 803019) was used for ICC. Alexa Fluor 488 conjugated to anti-β-amyloid 1-16 6E10 (1:200) (Biolegend, 803001) with 488 Conjugation kit (1:200) (Abcam, ab236553) performed per manufacturer instructions was used for live imaging.

### MetLuc ELISA

The MetLuc reporter assay was performed using Ready-To-Glow™ Secreted Luciferase Reporter Assay (Clontech, 631728) per the manufacturer’s instructions in a white 96-well (flat-bottom) plate (Greiner Bio-One, 655074). Luciferase activity was measured using the BioTek Cytation3 multi-mode plate reader. Luminescence readings were recorded as relative light units (RLU).

### Image analysis and statistical analysis

Videos and images were generated using ZEISS ZEN Microscopy Software (RRID:SCR_013672) and IMARIS software (RRID:SCR_007370). Image processing was done with IMARIS. Each tile scan comprised a total area of approximately 800 µm × 800 µm. Regions of interest (ROI) were generated using the surface function. ROI were generated based on the AβO-FITC channel, and the same settings were used for all treatment conditions replicate by replicate. FITC and chLecanemab mean pixel and sum pixel intensities within AβO-FITC ROIs were exported into IBM SPSS Statistics for statistical analysis. Assumption tests were performed for all analyses. Box plots were used to assess outliers, normality was assessed with the Shapiro-Wilk test (*p* > .05), and homogeneity of variances with Levene’s test (*p* > .05). ANOVA and Tukey multiple comparisons were performed when ANOVA assumptions were met. Welch’s *F* test and Games-Howell multiple comparisons were performed when the assumption of homogeneity of variances was violated. Descriptive statistics include means and standard deviations. All pairwise comparisons were two-tailed. Exact *p*-values are reported for significant and non-significant results.

