## Supplementary Figures for "Aβ-targeting synNotch Receptor for Alzheimer’s Disease: Expanding Applications to Extracellular Protein Aggregates"

**Supplementary Materials.**

***Video 1.*** *Time-lapse video of confocal tile scan images of untransduced NIH 3T3 cells that do not produce chAducanumab in response to Aβ treatment. Time-lapse images are taken at 20x magnification, starting 2 hours after addition of AβO 0.1 µM plus AβF 0.5 µM as well as anti-mouse IgG2a secondary antibody to identify chAducanumab Fab-Fc.*

***Video 2.*** *Time-lapse video of confocal tile scan images of Adu-synNotch-expressing NIH 3T3 cells producing chAducanumab in response to Aβ treatment. Time-lapse images are taken at 20x magnification, starting 2 hours after addition of AβO 0.1 µM plus AβF 0.5 µM as well as anti-mouse IgG2a secondary antibody to identify chAducanumab Fab-Fc.*

***Video 3.*** *Time-lapse video of confocal tile scan images of untransduced NIH 3T3 cells treated with Aβ-positive 6E10 aggregates. Time-lapse images are taken at 20x magnification, starting 6 hours after addition of AβO 0.1 µM plus AβF 0.5 µM as well as 6E10-AF488 conjugated antibody to identify Aβ.*

***Video 4.*** *Time-lapse video of confocal tile scan images of untransduced NIH 3T3 cells that do not produce chAducanumab in response to Aβ treatment. Time-lapse images are taken at 20x magnification, starting 6 hours after addition of AβO 0.1 µM plus AβF 0.5 µM as well as anti-mouse IgG2a secondary antibody to identify chAducanumab Fab-Fc.*

***Video 5.*** *Time-lapse video of confocal tile scan images of Adu-synNotch-expressing NIH 3T3 cells treated with Aβ-positive 6E10 aggregates. Time-lapse images are taken at 20x magnification, starting 6 hours after addition of AβO 0.1 µM plus AβF 0.5 µM as well as 6E10-AF488 conjugated antibody to identify Aβ.*

***Video 6.*** *Time-lapse video of confocal tile scan images of Adu-synNotch-expressing NIH 3T3 cells producing chAducanumab in response to Aβ treatment. Time-lapse images are taken at 20x magnification, starting 6 hours after addition of AβO 0.1 µM plus AβF 0.5 µM as well as anti-mouse IgG2a secondary antibody to identify chAducanumab Fab-Fc.*

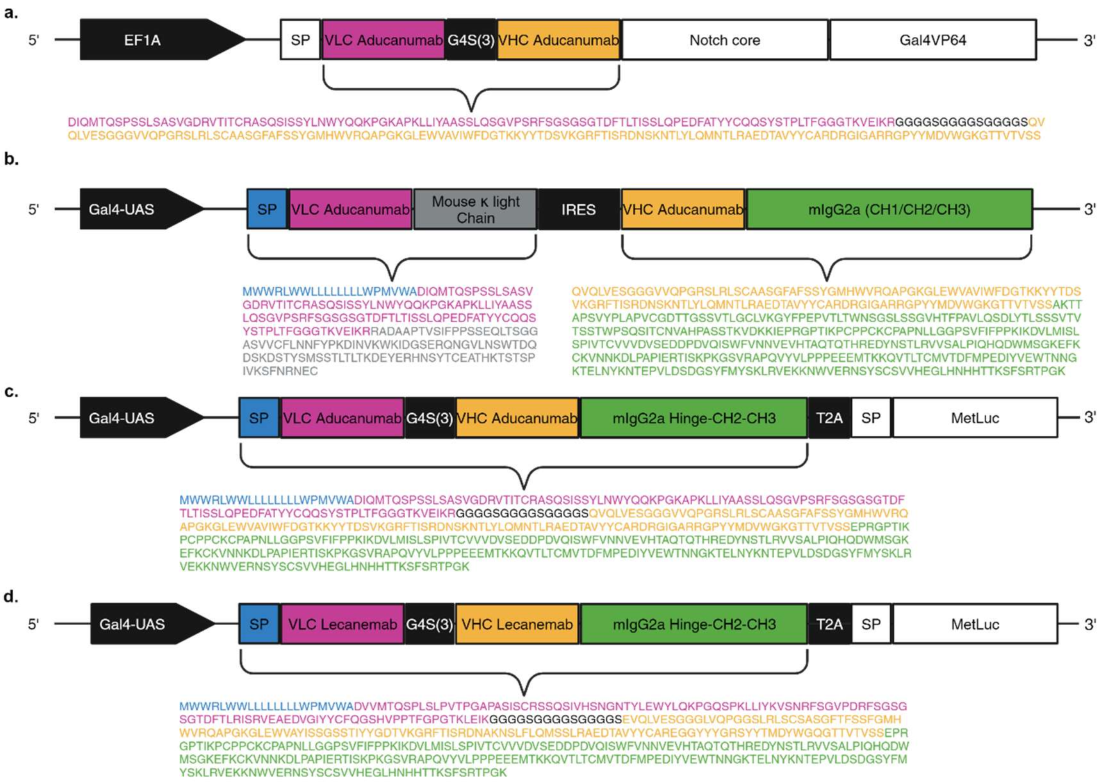


***Supplementary Figure 1. Amino acid sequences for relevant constructs used in the experiments. (a)*** *Sequence for the scFv of Adu-synNotch. (**b**) Sequence for chAducanumab in standard antibody format (Fab-Fc). (**c**) Single chain chAducanumab followed by an in-frame MetLuc reporter separated by T2A sequence with an N-terminal added Furin cleavage site and a GSG linker. (**d**) Single chain chLecanemab followed by an in-frame MetLuc reporter separated by T2A with a Furin cleavage site and a GSG linker as in (**c**).*

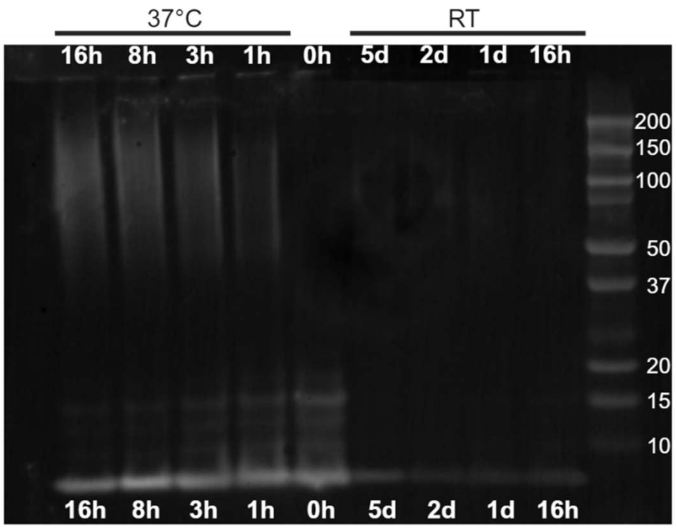


**Supplementary Figure 2. Aβ$_{(1-42)}$ aggregation time course shows increasing MW with incubation time at 37°C.** *Western blot of 6E10-stained Aβ$_{(1-42)}$ preparations 0-16 hours at 37°C or 16 hours to 5 days at room temperature (RT). Lower bands are consistent with the MW of Aβ monomers. At 0 hours, there are already higher MW species, which decline with incubation times, while higher MW species accumulate when incubated at 37°C but not at RT. Incubation times of 7 days at 37°C further enrich higher molecular weight (data not shown).*

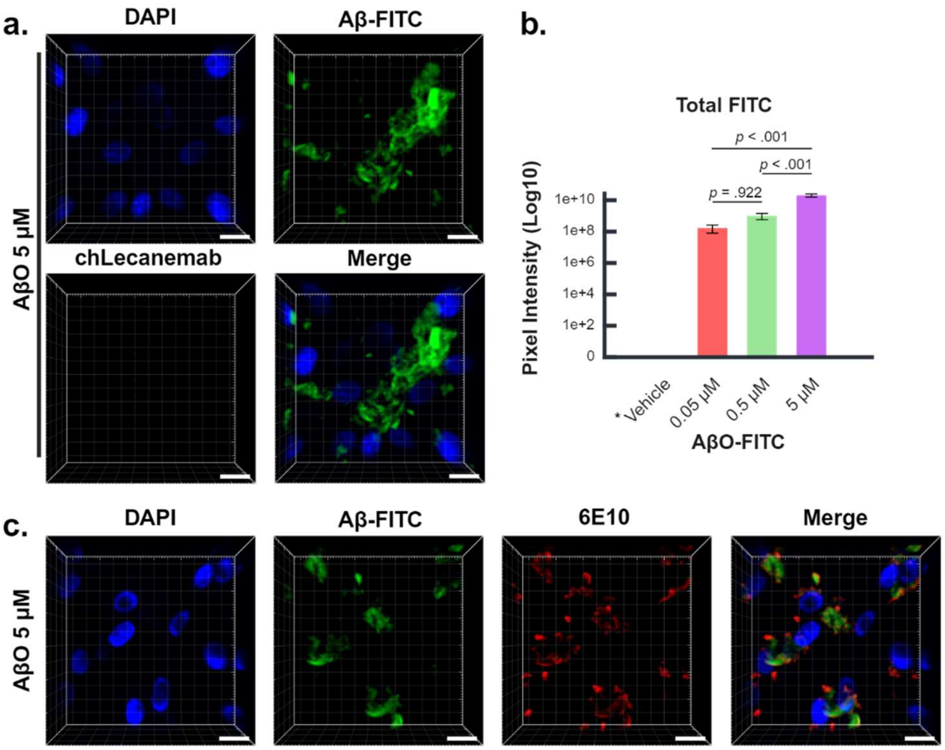


***Supplementary Figure 3. (a)*** *Representative 3D confocal images of 5 μM AβO-FITC treated untransduced NIH 3T3 cell controls of experiments described in Figure 3. Immunostaining for chLecanemab with mouse IgG2a secondary antibody. Aβ aggregates are identified by a FITC tag and cell nuclei by DAPI. chLecanemab signal was absent from Aβ-FITC aggregates. Representative images of three independent experiments.* ***(b)*** *Quantification of total (sum) FITC signal (Log 10 scale for easy visualization). Multiple comparisons with Tukey's (HSD) test performed on untransformed pixel intensity values. *Vehicle was not quantified due to lack of detectable FITC signal. Results are from three independent experiments.* ***(c)*** *Representative 3D confocal images of 5 μM AβO-FITC treated untransduced NIH 3T3 cells immunostained with pan-Aβ antibody 6E10 show colocalization with FITC. Error Bars 1x SE. Scale bar: 20 μm.*
